# A Supervised Contrastive Framework for Learning Disentangled Representations of Cell Perturbation Data

**DOI:** 10.1101/2024.01.05.574421

**Authors:** Xinming Tu, Jan-Christian Hütter, Zitong Jerry Wang, Takamasa Kudo, Aviv Regev, Romain Lopez

**Affiliations:** University of Washington; Genentech Research and Early Development; California Institute of Technology; Stanford University

## Abstract

CRISPR technology, combined with single-cell RNA-Seq, has opened the way to large scale pooled perturbation screens, allowing more systematic interrogations of gene functions in cells at scale. However, such Perturb-seq data poses many analysis challenges, due to its high-dimensionality, high level of technical noise, and variable Cas9 efficiency. The single-cell nature of the data also poses its own challenges, as we observe the heterogeneity of phenotypes in the unperturbed cells, along with the effect of the perturbations. All in all, these characteristics make it difficult to discern subtler effects. Existing tools, like mixscape and ContrastiveVI, provide partial solutions, but may oversimplify biological dynamics, or have low power to characterize perturbations with a smaller effect size. Here, we address these limitations by introducing the Supervised Contrastive Variational Autoencoder (SC-VAE). SC-VAE integrates guide RNA identity with gene expression data, ensuring a more discriminative analysis, and adopts the Hilbert-Schmidt Independence Criterion as a way to achieve disentangled representations, separating the heterogeneity in the control population from the effect of the perturbations. Evaluation on large-scale data sets highlights SC-VAE’s superior sensitivity in identifying perturbation effects compared to ContrastiveVI, scVI and PCA. The perturbation embeddings better reflect known protein complexes (evaluated on CORUM), while its classifier offers promise in identifying assignment errors and cells escaping the perturbation phenotype. SC-VAE is readily applicable across diverse perturbation data sets.

## 1 Introduction

The advent of CRISPR technology has revolutionized functional genomics, thereby enabling precise and targeted genetic manipulations [1]. The adoption of single-cell RNA-Seq (scRNA-Seq) as a readout (Perturb-Seq) has further advanced this field by combining a high content readout with high-throughput capabilities and single-cell resolution [2, 3]. These technologies overcome the limitations of traditional pooled screens, which were limited to simpler readouts such as cell death or marker protein expression, and of arrayed screens with bulk RNA-seq, which had limited scale and only bulk resolution. The high resolution, massive scale nature helped reveal the impact of gene perturbations on cell-to-cell variations that are critical for understanding complex biological phenomena [4], from immune cell activation [3] to cancer cell-intrinsic resistance mechanisms [5]. While initial screens focused on the impact of dozens or hundreds of perturbations, they have since grown to genome-scale screens that drastically improved the efficiency and depth of functional genomics [6].

The computational analysis of Perturb-Seq screens comes, however, with significant challenges. First, the scRNA-seq measurements are high-dimensional, and affected by strong technical noise [7]. Some sources of technical noise are those common to all scRNA-seq studies, such as the sequencing depth and count over-dispersion, but others are specific to the perturbation context. For example, the capture rate of the guide RNA may be noisy, and therefore the association of a single cell to a perturbation may have some error [8]. Second, guide expression in a cell does not not necessarily mean that the cell has been genetically perturbation, because of varying efficiency rates of the Cas9 enzyme and the time delay from genetic perturbation to an effect on the target protein [9, 3]. Third, the perturbation effect size may be relatively low for most perturbations [3], making them more challenging to identify. This is especially the case, because the effect size is opften comparable to other sources of biological variation, such as the different stages of the cell cycle [5]. In this case, established approaches to single-cell analysis such as Principal Component Analysis (PCA) or generative models [10] may not capture the effect of perturbations, as they tend to focus on patterns with highest variance across the whole data set (e.g., stages of the cell cycle).

Researchers introduced methods to address all or some of these challenges. For example, both MIMOSCA [3] and Mixscape [11] identifies cells that carry a guide but do not exhibit the impact of a genetic peturbation (denoted “escaping” cells below). Both approaches first treat the inferred assignment of cells to perturbation as ground truth, uses it to identify a perturbation-specific signal, and then reassesses the cells in light of this model to identify the escaping cells as the ones who more closely resemble unperturbed than perturbed cells (despite their guide assignment). Mixscape further uses linear discriminant analysis (LDA) to visualize the perturbed cells. However, MIMOSCA’s linear model and Mixscape’s use of LDA and a two-component Gaussian mixture model likely oversimplify real-world complexities and fail to capture non-linear dynamics.

More recently, ContrastiveVI [12] proposed to analyze Perturb-Seq data by learning two latent spaces: a *background* space that captures cell-to-cell variation that is present in the non-perturbed population, and a *salient* space that encodes variation from the genetic perturbations. This model is based on the Contrastive Analysis (CA) framework [13, 14] as well as the single-cell Variational Inference [15] (scVI) deep generative model. Although the learning of such *disentangled* representations is an exciting prospect, this objective is itself quite challenging, as it requires careful model design and regularization [16, 17]. Indeed, ContrastiveVI employs a regularization introduced in [18] to avoid leakage of the background into the salient space. However, it is not agreed upon which regularization works best, and there may therefore exist challenging data sets on which ContrastiveVI’s performance is sub-optimal. Additionally, ContrastiveVI assumes that only two data sets are being processed, both stemming from identically and independently distributed data distributions: the background distribution and the target distribution. Therefore, the estimates of the latent variables from the salient space are shrunk towards the prior, potentially resulting in a lower power for discerning small sized effects. Finally, ContrastiveVI does not propose a mechanism for identifying and removing cells that have escaped CRISPR perturbation.

Here, we address these limitations with Supervised Contrastive Variational Autoencoder (SC-VAE), a novel framework for learning disentangled representations from Perturb-Seq data. Following the CA framework, SC-VAE learns two latent spaces with the same semantic, but also jointly models guide RNA identity alongside gene expression measurements. Implementation-wise, this amounts to using the salient space to predict the assignment of cells to perturbations. The idea of biasing a representation to be discriminative of external labels is quite common in the literature of topic models [19], but, to the best of our knowledge, has not been proposed in the context of CA. To address the problem of learning disentangled representations, we employed the Hilbert-Schmidt Independence Criterion [20] as a regularization technique, and demonstrated satisfactory results. We validated the performance of SC-VAE on two recent large-scale Perturb-Seq data sets [6]. First, we demonstrated that SC-VAE is more sensitive than ContrastiveVI and PCA, while quantifying the effect of a perturbation, suggesting that SC-VAE could improve over ContrastiveVI while characterizing perturbations with small effect sizes. Second, we showed that the salient embeddings from SC-VAE recapitulates known protein complexes with higher accuracy than those from PCA, scVI, and ContrastiveVI. Finally, we inspected the ability of the classifier to identify potential errors in the assignments of cells to perturbations. Our results suggest that SC-VAE’s classifier may be used to identify both false negative cells (perturbed, but without a detected guide), as well as escaping cells. Our approach is readily applicable to other perturbation data sets.

## 2 Methods

We developed SC-VAE to exploit the perturbation label (e.g., guide RNA identity) to better disentangle perturbation effects from natural cell-to-cell variations in large-scale Perturb-Seq. SC-VAE extends the CA framework by adding a supervision component to the generative model. Our framework is based on an auto-encoder architecture (Figure 1), and builds on recent work on CA [14, 18, 12].

**Figure 1:**
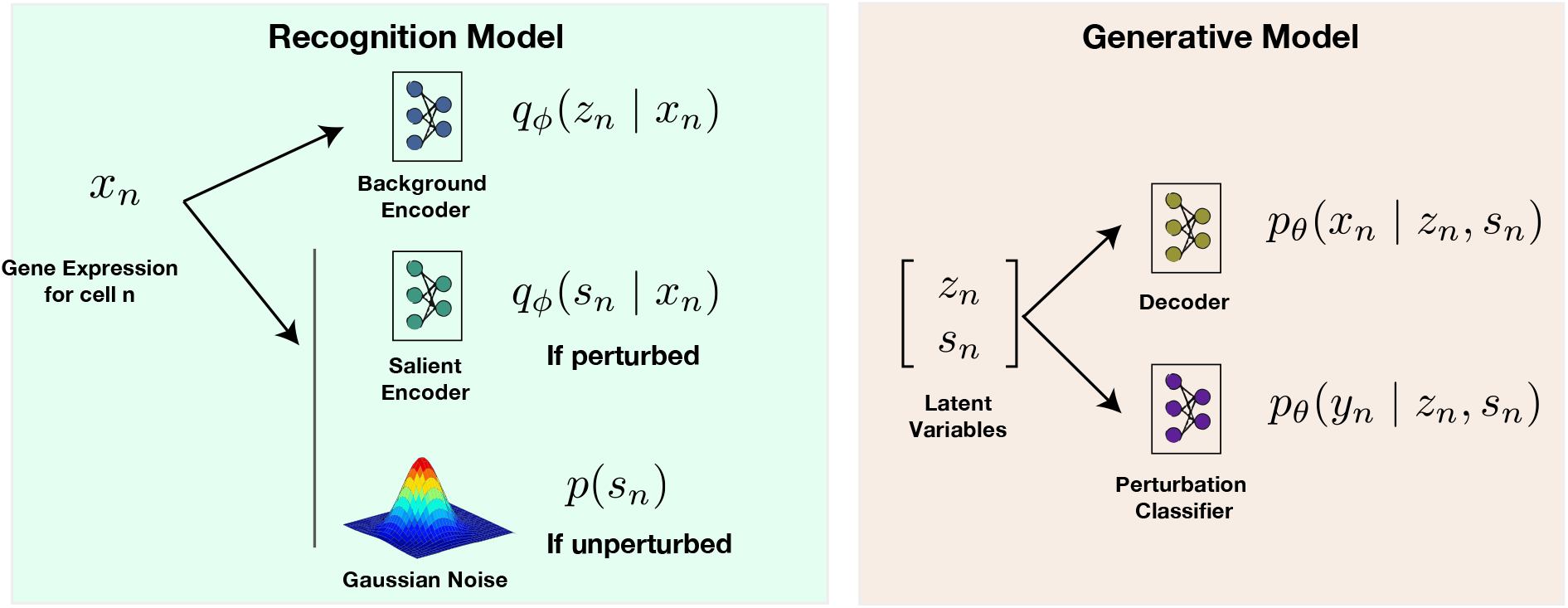
Model schematics: Our method incorporates two distinct encoders: a background encoder, capturing biological attributes like cell cycle processes, and a salient encoder, specifically targeting perturbation effects. Both latent embeddings are fed into a decoder that outputs the likelihood parameters for modeling gene expression levels. A perturbation classifier discerns perturbation labels based on the salient embedding.

### 2.1 Generative Model

For each cell *n*, let latent variable

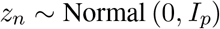

be a low-dimensional random vector encoding cell variations present in the control cells (the background data set). We refer to *z* as the *background* latent space, or embedding. Then, let latent variable

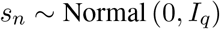

be a low-dimensional vector encoding variations due to perturbations. We refer to *s* as the *salient* latent space, or embedding. For each single cell *n* and gene *g*, the gene expression *x*_*ng*_ is generated as

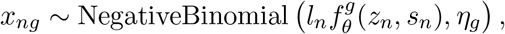

where *f*_*θ*_ is parameterized by a neural network, *l*_*n*_ is the number of RNA transcript molecules (reads) captured in cell *n* (referred to as library size, *l*_*n*_ =∑ _*g*_ *x*_*ng*_), and *η*_*g*_ are inverse-dispersion parameters. The modeling choices for this conditional distribution are identical to the ones from scVI [15], and ContrastiveVI [12]. Finally, we observe the perturbation identity *y*_*n*_ ∈ {∅, 1, …, *K*}, where ∅ denotes a non-targeting control (NTC) perturbation (i.e., no effect) and *K* denotes the number of distinct perturbations:

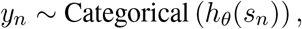

where the function *h* is a classifier parameterized by a neural network. We note that *h* takes as input only *s*_*n*_, which makes the two latent spaces asymmetric.

### 2.2 Variational Inference

The marginal probability of the data *p*_*θ*_(*x, y*) is intractable. We therefore proceed to posterior approximation with variational inference to learn the model’s parameters. Although the definition of the classifier breaks symmetry between the two latent spaces, this is not enough to prevent leakage between them in practice, and therefore we treat the cases *y*_*n*_ = ∅ and *y*_*n*_ ≠ ∅ differently. This helps constrain the inference towards learning disentangled representations.

For the perturbed cells (i.e., *y*_*n*_ ≠ ∅), we proceed following previous work [14, 12], and use a mean-field variational distribution *q*_*ϕ*_(*z*_*n*_ | *x*_*n*_)*q*_*ϕ*_(*s*_*n*_ | *x*_*n*_), where, as in the VAE framework [21], each *q*_*ϕ*_(*z* | *x*) and *q*_*ϕ*_(*s*| *x*) follows a Gaussian distribution with a diagonal covariance matrix. The evidence lower bound is derived as:

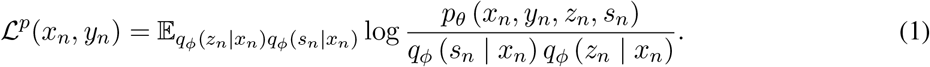

For the unperturbed cells, we want to render the latent variable *s*_*n*_ uninformative, so that only *z*_*n*_ is used to model the expression levels. Therefore, we derive a lower bound such that the variational distribution for latent variable *s*_*n*_ is ignored for the reconstruction error of *x*_*n*_ (but not *y*_*n*_):

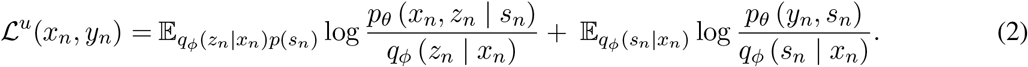

The derivations appear in Appendix A. To optimize this lower bound, we proceed exactly as in stochastic variational inference, by subsampling at each iteration a mini-batch of cells from the training set, and for each cell, taking one sample *z*_*n*_ and *s*_*n*_ from the variational distribution using the reparameterization trick, and one sample *s*_*n*_ from the prior distribution *p*(*s*_*n*_).

### 2.3 Regularization Strategy

We regularize the model by employing the Hilbert-Schmidt Independence Criterion (HSIC) [20] to further disentangle the latent space. This technique builds upon the framework of HSIC-constrained Variational Auto-Encoders (HCVs) [16] to enforce statistical independence between the background and salient latent variables. Throughout all the experiments, we weighted the HSIC term by the same scalar that was adjusted so that the ELBO and the HSIC have similar magnitude during training.

## 3 Results

We demonstrate the performance of SC-VAE on two large Perturb-Seq data sets [6]: one from thelymphoblast K562 cell line with knock-down perturbation (CRISPRi) of 2,058 essential genes, and the second from the RPE1 immortalized epithelial cell line with knock-down perturbation of 2,394 essential genes.

### 3.1 Visualization of Learned Latent Representations

We first investigated the result of SC-VAE applied to the K562 cell line CRISPRi data set. To better visualize data from diverse perturbation groups, we focused on the top 100 perturbations based on number of cells (i.e., those leading to most cell proliferation), and performed two exploratory analyses. In one, we confirmed that the classifier did not cause the latent variable model to overfit and separate cells from different perturbation labels in a perfect and arbitrary fashion. If this occurred, the model would even perfectly delineate labels from perturbations with no discernible effect, but only on the training data. As a simple diagnosis, we split our data into a training and a test set, and embedded cells from the test set alongside the training set, for both the salient space and the background space (Figure 2, left). The distribution of points across data splits is visually balanced across both spaces, indicating that the model is not blatantly overfitting. In the other analysis, we colored cells according to their perturbation labels *y*_*n*_ in both latent spaces. While many cells clustered together by perturbations in the salient space, such patterns are barely noticeable in the background space (Figure 2, right), suggesting that the method behaves as expected.

**Figure 2:**
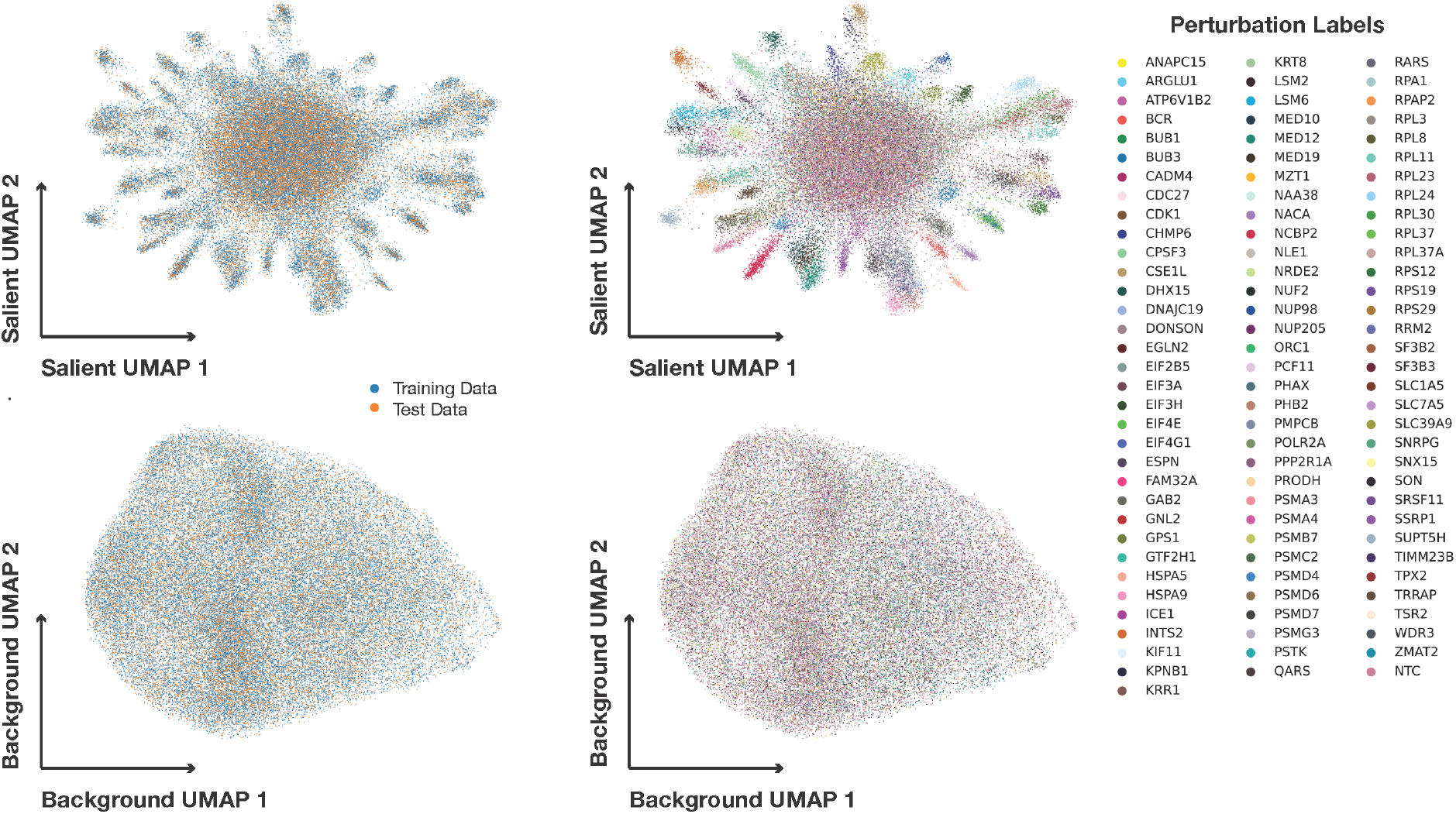
UMAP visualization of cell profiles (dots) in the background (bottom) and salient (top) spaces inferred by SC-VAE applied to the top 100 perturbations from the K562 Perturb-seq data set, with cells colored by data split (left) or perturbation label (right).

Next, we analyzed the full RPE1 Perturb-seq data set, and visually compared its results to PCA as a simple baseline (more thorough benchmarking is presented below). For visualization purposes, we color coded the 2,394 essential gene perturbations by eight prominent categories annotated in the original study [6]: exosome, mediator complex, mitchochondrial protein translocation, nucleotide excision repair, and ribosomal units S39, S40 and S60. We categorized the remainder as “others”, while also considering the control cells as a separate group. Again, visual inspection confirmed that the SC-VAE model did not not overfit (Figure 3A, left).

**Figure 3:**
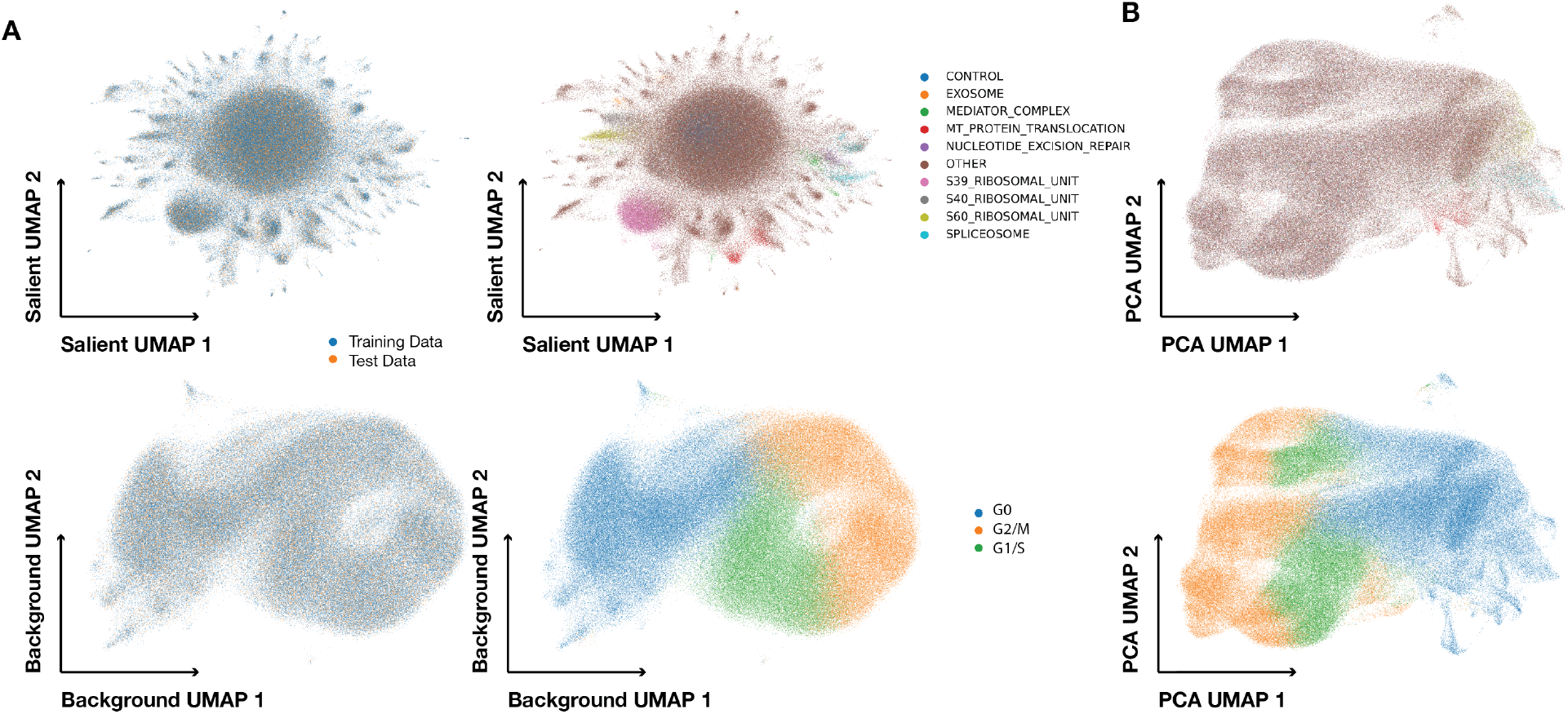
UMAP visualization of the background space and the salient space inferred by SC-VAE applied the RPE1 Perturb-seq data set. (A, left) Cells are colored according to the data split. (A, right) Cells are colored according to their perturbation group for the salient space, and by cell cycle phase for the background space. (B) Same results for PCA.

While the SC-VAE salient space showed cells forming multiple small clusters, many with distinct annotations, PCA data showed much less variation attributable to the genetic perturbations (Figure 3B). We hypothesized that PCA was mainly accounting for differences within the unperturbed cells, as previously documented [12]. Indeed, when annotating profiles by the scores of cell cycle phase signatures[22] (via scanpy), cell cycle phase was the main cell-to-cell heterogeneity present in the PCA space, as well as in the SC-VAE background (but not salient) space.

These results suggest that SC-VAE unveils non-trivial variations in perturbation data sets, and groups perturbations according to their biological function in the salient space, while separately capturing biological variation from the unperturbed cells (e.g., cell cycle) in the background space. We now turn to rigorous benchmarking to strengthen these claims.

### 3.2 Univariate benchmarks

As a first benchmark, we aimed to quantify the ability of different latent spaces to distinguish perturbed from unperturbed cells (univariate benchmarks per [23]). This allows us to compare the sensitivity of our salient space to other methods, and how well we learned a disentangled representation, such that perturbed cells are close to control cells in the background space (except if the perturbation affects biological variation occurring in the background, such as cell cycle). To this end, we relied on the energy distance, following [6], a metric to compare two distributions based on the pairwise distances between their samples. Let *X* and *Y* be two random variables, and 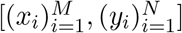 be some independent draw from these random variables. The empirical estimate *Ê* (*X, Y*) of of the energy distance between these two distributions, *E*(*X, Y*), accounting for the pairwise distances within and between the sets, is derived as:

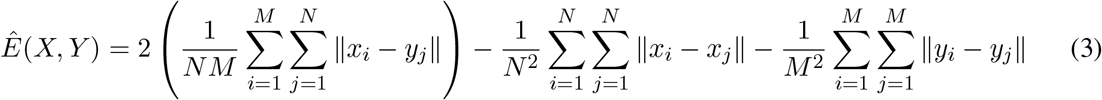

The first term in *Ê* (*X, Y*) computes the mean distance between points in sets *X* and *Y*, while the second and third terms gauge the internal pairwise distances within *X* and *Y* . If *E*(*X, Y*) = 0, the random variables are identically distributed; a higher *E*(*X, Y*) indicates greater distributional divergence between random variables *X* and *Y* .

We calculated the pairwise energy distance between each of the top 100 perturbation labels and the control population, using both latent spaces of SC-VAE and ContrastiveVI, and in PCA space. To mitigate the impact of the difference in scale between latent spaces, we used the cosine distance in place of the Euclidean norm in (3). We first compared the energy distance matrices between the background and the salient space of SC-VAE (Figure 4A). As expected, the salient space induces a much higher energy distance compared to the background space, suggesting that the two spaces are disentangled. Moreover, the energy distances for SC-VAE’s salient space were consistently higher than those for ContrastiveVI’s salient space or for the PCA space (Figure 4B, top). This suggests that our salient space is much more sensitive than the one from other methods. Conversely, the energy distances for SC-VAE’s background space were consistently lower than ContrastiveVI’s background space and the PCA space (Figure 4B, bottom). This suggests that our background space contains less information about the perturbation compared to other methods, and therefore that our latent spaces are better disentangled.

**Figure 4:**
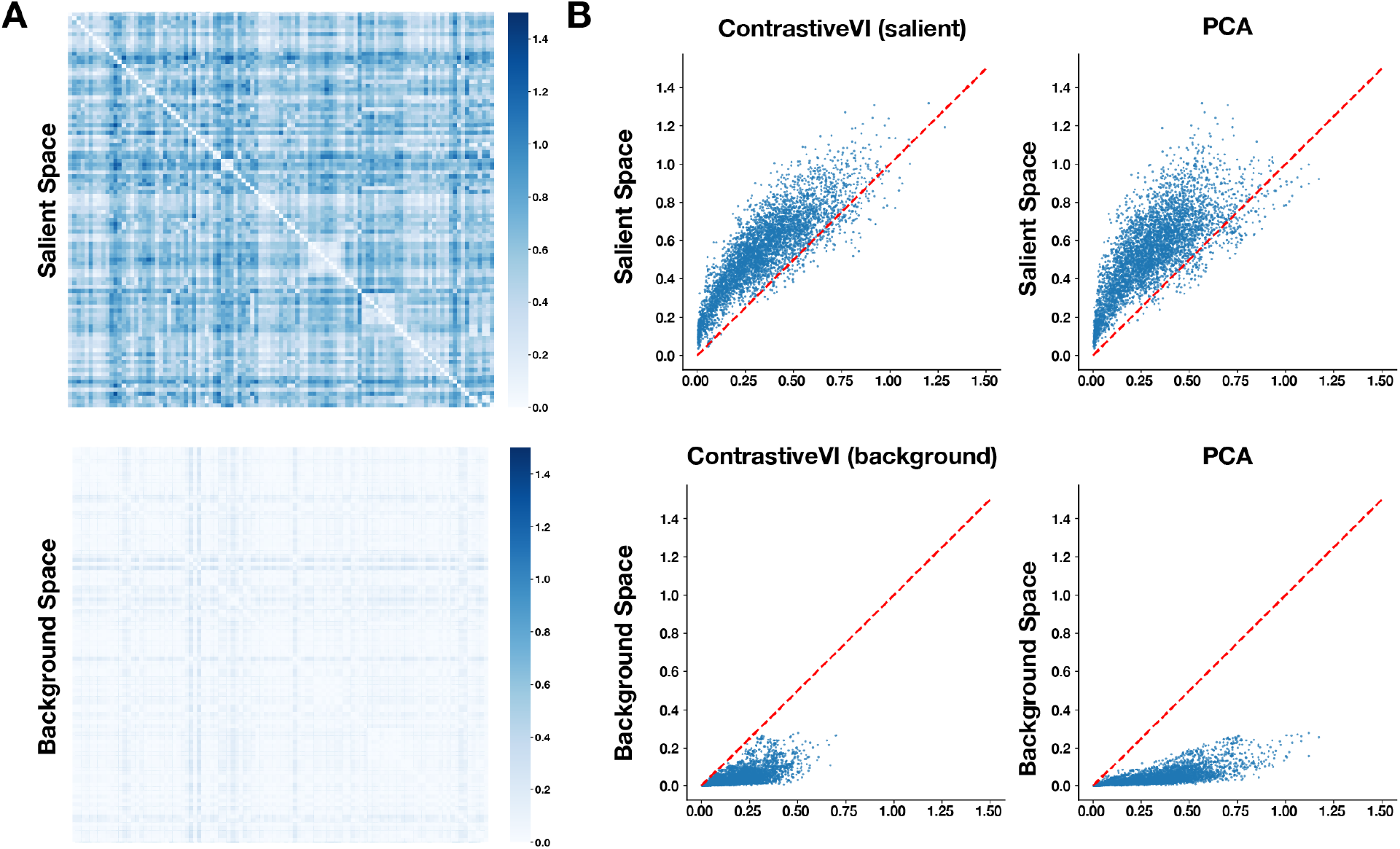
Comparison of the energy distance values. (A) Comparison of the distance matrices for all perturbation groups, calculated on the salient (top) and background (bottom) space of SC-VAE. (B) Benchmarking of SC-VAE compared to ContrastiveVI (left), and PCA (right). Baseline (x-axis) and SC-VAE (y-axis) energy distance values for each pair of perturbations (point) in the salient (top) and background (bottom) space.

### 3.3 Multivariate benchmarks

We next asked if SC-VAE can capture how perturbations are related to each other such that learned perturbation map is biologically relevant [23]. To test this, we relied on the well-established assumption that perturbations in genes encoding members of the same protein complex should lead to similar phenotypes an thus more similar cell profiles [6, 8]. Accordingly, We adapted previous methodology, using the CORUM complex database [24] and the following process:

- **Cell Aggregation**: We grouped cell profiles by perturbation, and calculated the mean embedding of all cells in each group as the aggregated embedding.
- **Shift Calculation**: We obtained the shift vector by subtracting the embedding of the control group from each perturbation embedding.
- **Similarity & Recovery**: We computed the cosine similarities between perturbations, and then predicted gene-gene relationships through varying cut-off thresholds on pairwise cosine similarity. We compared these with ground truth gene-gene relationships defined by co-membership in the same CORUM cluster.
- **Performance Evaluation**: We used a precision-recall curve analysis to evaluate the accuracy of our derived relationships against the complex database.

We compared the results of SC-VAE with ContrastiveVI, PCA and scVI (Figure 5). First, the salient embedding of SC-VAE has the best performance on this benchmark, outperforming the salient space of ContrastiveVI, but also the latent spaces from PCA and scVI. Second, the background space of SC-VAE has the lowest performance (followed by the background space of ContrastiveVI). This suggests that both methods are able to disentangle the spaces (at least partially), but SC-VAE might do so more effectively.

**Figure 5:**
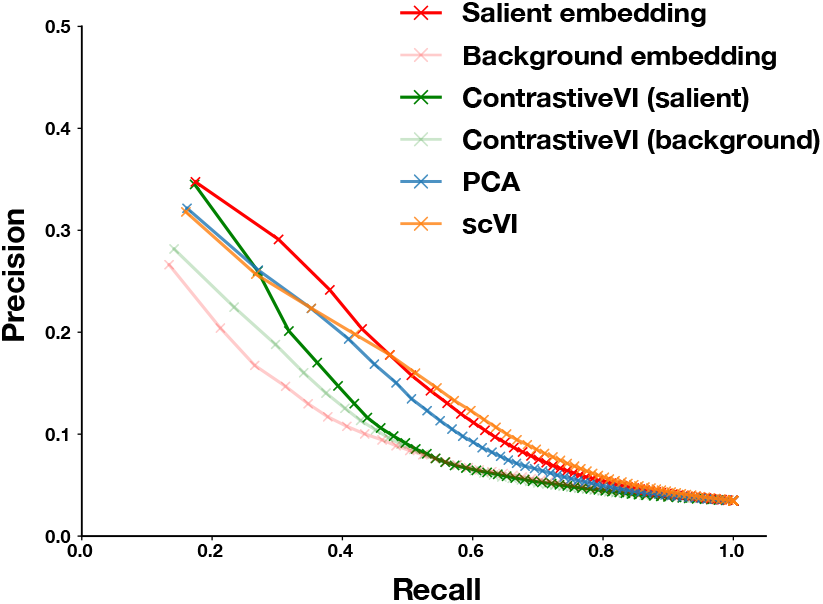
Precision-recall curves for the multivariate benchmark for SC-VAE, ContrastiveVI (both salient and latent), scVI and PCA. All four embedding spaces have the same dimension.

### 3.4 SC-VAE’s classifier predictions identified escaped and false negative cells

Integrating a classifier within the contrastive analysis framework presents an important practical advantage. It is possible to get predictions for the perturbation assignment of cells in a test set^1^. We therefore classified cells in the test set using the neural network *h*_*θ*_ applied to the salient embedding values, and reported the confusion matrix, comparing the transcriptome-inferred labels with the experimentally-inferred perturbation labels.

A close examination of the confusion matrix, displayed in logarithmic scale, reveals three distinct patterns (Figure 6A). First, the diagonal line indicates that the classification is overall accurate. Second, a vertical line shows numerous cells from varied perturbation groups classified as control cells. We posit that these are in fact escaping cells. Third, some control cells (with an NTC barcode detected) are categorized under other perturbation groups, suggesting the presence of cells falsely classified as negative control cells, or negative control cells that exhibit perturbation phenotypes due to their underlying heterogeneity.

**Figure 6:**
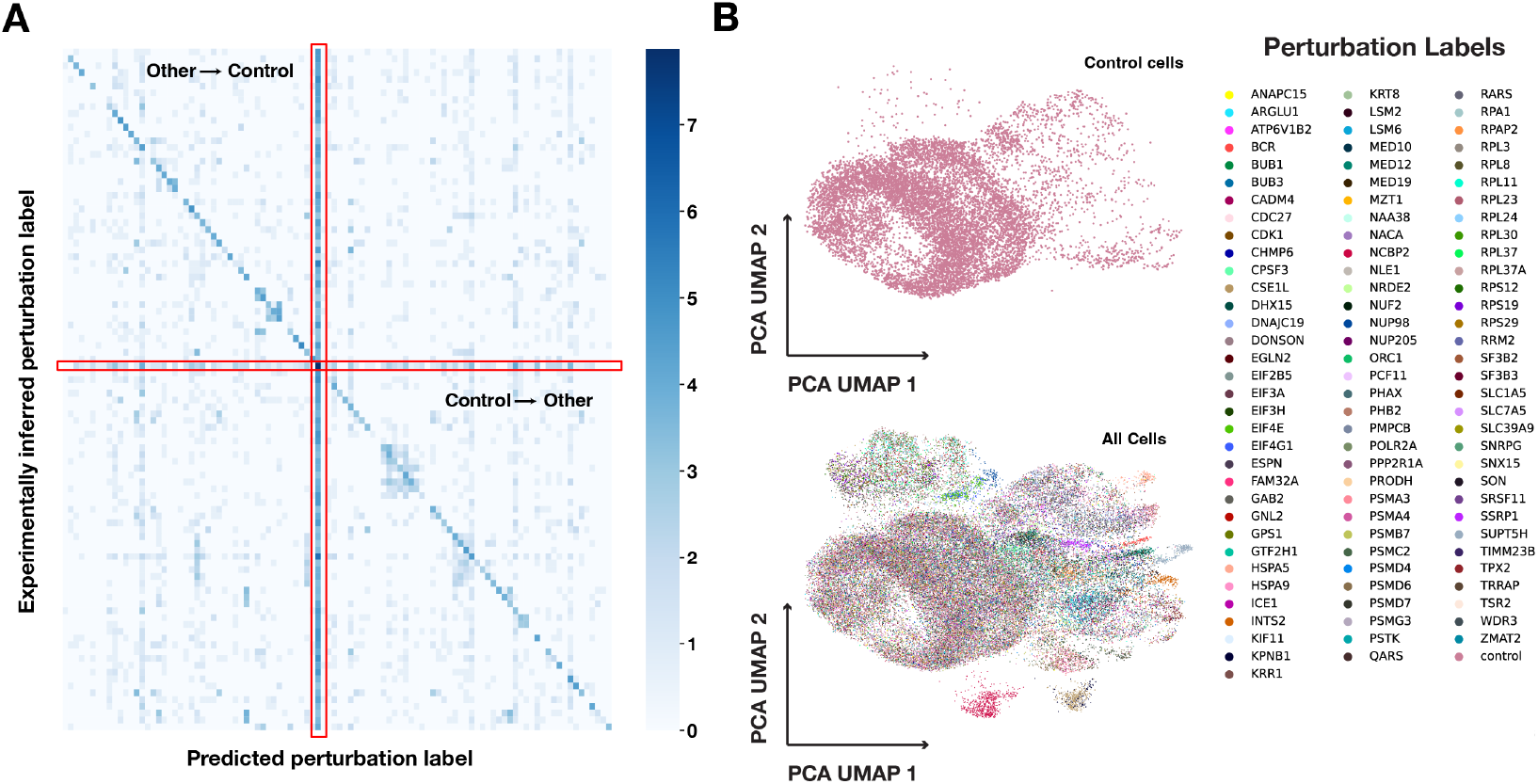
Visualization of SC-VAE’s classifier output. (A) Confusion matrix of predicting perturbation labels, calculated on a test set and colored according to a logarithmic scale. The red rectangles delineate the position of the label that corresponds to the unperturbed (control) cells. (B) Position of the control cells in the UMAP visualization of the space of principal components. Many control cells lie alongside other perturbations.

To substantiate these observations, we further assessed the distribution of control cells in an unsupervised space using PCA (calculated on the complete data set) and visualized it via UMAP (Figure 6B). Notably, a minor fraction of the control cells indeed clustered with perturbed cells, especially in the upper right segment of the visualization.

## 4 Conclusion

In this study, we introduced SC-VAE, a novel computational framework, a supervised contrastive method for learning disentangled representations of cell perturbation data, for analyzing Perturb-seq data. SC-VAE outperformed existing methods in terms of both accuracy and interpretability. One of the key advantages of the SC-VAE framework is its ability to disentangle the effects of perturbations from the background cellular context. This is achieved by learning a latent space that separates the perturbation effects from the background, which allows for more accurate and interpretable data analysis.

## Acknowledgments and Disclosure of Funding

We thank Rebecca Boiarsky, Antonio Rios, and Kexin Huang for insightful feedback throughout the duration of this project which greatly improved this work. We also thank members of the Regev Lab and the Research Biology Artificial Intelligence department (BRAID) at Genentech for providing constructive feedback on the results presented in this work.

## A Evidence lower bound for unperturbed cells

We provide here the derivation for the evidence lower bound we implemented for unperturbed cells. In order to provide a separate treatment of the transcriptomic profiles *x*_*n*_ and the perturbation label *y*_*n*_ for these cells, we derive a lower bound on the composite likelihood:

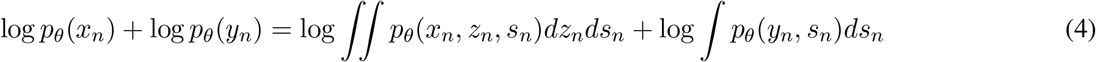

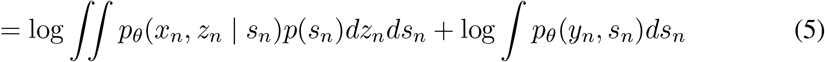

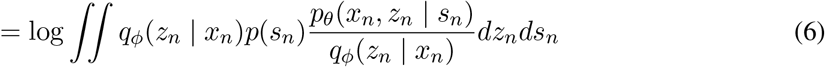

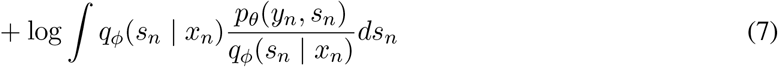

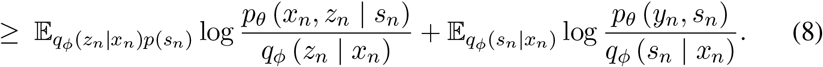

## Footnotes

1 In principle, it is also possible to get predictions for cells in the train set, but we advise against this: even if the model does not show signs of overfitting on average, the predictions will understandably be biased towards the value in the training set.

